# Incidence and multidrug resistance of *Escherichia coli* pathotypes on fresh vegetables and salads

**DOI:** 10.1101/2024.07.18.604175

**Authors:** C. R. Vázquez-Quiñones, M.A Rincón-Guevara, I. Natividad-Bonifacio, C. Vázquez-Salinas, H. González-Márquez

## Abstract

Diarrhea is a severe health problem and one of the leading causes of child mortality in Africa, Asia, and Latin America. Diarrhea is often caused by consuming contaminated food and improperly washed fruit and vegetables that harbor norovirus, *Campylobacter*, non-typhoid *Salmonella*, and pathogenic *Escherichia coli.* The research examined 334 samples of ready-made salads with lettuce, carrots, tomatoes and fresh coriander and lettuce. Genotyping involved detecting *st* and *lt* genes for enterotoxigenic *E. coli* (ETEC) and the *uid*A gene for beta-glucuronidase. ETEC was found in 51.56% of confirmed isolates, accounting for 9.9% of samples gathered in two years. Contamination rates by ETEC were 78.78% in coriander, 9.09% in lettuce, 9.09%, and 3.03% in green salads from La Vicentina and La Purísima markets, respectively. Among samples, 12.12% have both fragments (*st/lt*), 33.3% have only the *lt* fragment, and 54.6% have just *st*. In salads, the *lt/st* genes were detected in 9.09% (3), *lt* in 3.3% (1), and *st* was absent. In coriander, 21.21% have *lt*, 51.51% *st*, and 6.06% showed amplification for both. ETEC was found in 9.09% of the confirmed lettuce strains, with 3.03% *lt* gene, 3.03% *st* gene, and 3.03% both (*lt/st*). There are no reported data with the same ratios for Mexico City. ETEC’s presence in products consumed in markets or streets in Mexico City, coupled with lifestyle changes that have increased vegetable consumption, is a significant concern. These findings underscore the potential health implications and the urgent need for further investigation and preventive measures.

## 2 Introduction

Diarrheal diseases are a severe health problem and one of the leading causes of child mortality, particularly in Africa, Asia, and Latin American countries (1). The World Health Organization estimates almost 1.7 billion episodes of diarrhea in children globally each year, resulting in the fatality of nearly 450,000 infants under the age of five and another 50,000 children between 5 and 9 years old (2).

Diarrheal diseases are the most common manifestation of foodborne illness, with children being especially vulnerable (3). In Mexico, intestinal illness represented the second leading cause of morbidity in children under 5 years old, according to the 2022 Annual of Morbidity from the Epidemiology General Management. Additionally, the Mexican General Directorate of Epidemiology bulletin reported over 600,000 cases of acute diarrheal illness (EDA) the following year, underscoring the public health significance of this issue (4). Diarrhea is often caused by consuming contaminated meat, raw or undercooked eggs, and improperly washed fruit and vegetables that harbor norovirus, *Campylobacter*, non-typhoid Salmonella, and pathogenic *Escherichia coli* (3).

*E. coli* is a common inhabitant of the mammalian digestive tract and is typically transmitted via the fecal-oral route. Diarrheagenic *E. coli* harbors virulence factors that enable its adherence to the intestinal mucosa, leading to persistent infections, inflammation, and lesions in the host and an immunologic reaction characteristic to each case. These characteristics have led to classification of *E. coli* into six pathotypes: Adherent invasive *E*. *coli* (AIEC); diffusely adherent *E. coli* (DAEC); enteroaggregative *E. coli* (EAEC); enteroinvasive *E. coli* (EIEC); enteropathogenic *E. coli* (EPEC); enterotoxigenic *E. coli* (ETEC) and Shiga toxin-producing *E. coli* (STEC) (5, 6).

Fruit, vegetables (especially fresh salads), and seasonings are among the foods that can be contaminated with one of these microorganisms. Over the past 30 years, global per capita consumption of fresh vegetables has increased by 25%, with production rising 30% in the same period (from 30 to 60 million metric tons). Vegetables are commonly consumed both in establishments and on the streets. It is well known that vegetable consumption is associated with a rising number of illnesses due to bacterial contamination and poor hygienic practices during processing (7).

From a health perspective, it is crucial to document better the prevalence of pathogens transmitted through vegetables. While the per capita consumption of fresh vegetables in the form of salads in Mexico is unknown, the mean global consumption is estimated at 186 g/day, with Central America at 56 g/day and Central Asia at up to 349 g/day (8).

Horticultural products can get contaminated at various points along their production chain, from crop growth conditions, including residual water usage for irrigation, to the point of sale and even during consumption. This contamination raises concerns about the increasing presence of pathogens such as *E. coli* producing broad-spectrum beta-lactamases (ESBL *E. coli*) (9–11). Antimicrobial resistance among diverse microorganisms is an escalating public health emergency due to the difficulties in treating these infections and the increasing prevalence of multi-antibiotic-resistant strains (12).

This study aims to identify the presence of *E. coli* pathotypes in fresh salads, lettuce, and coriander sold or consumed in markets and on the streets in some regions of Mexico City and to determine the prevalence of resistance to the most common antibiotics used for the treatment of gastrointestinal illness in Mexico.

## 3 Results

### 3.1 Microbiological analysis

From the 128 analyzed samples, 334 isolates with morphological characteristics consistent with *E. coli* were obtained. Subsequently, seventy-eight isolates were confirmed as *E. coli* through biochemical testing. These isolates were obtained from coriander samples (50%) and twenty-six from lettuce samples (33.33%). Ten isolates were obtained from fresh green salads containing cucumber and carrot (EV1) (12.82%), while only three isolates were obtained from the salads containing cucumber, carrot, lettuce, and beetroot (3.85%) (Table 1).

**Table 1.** Isolated strains were identified and confirmed by PCR and MALDI TOF as *Escherichia coli* isolated from vegetables and salads in Mexico City.

| Sample<br>(type) | Sample<br>(number) | *E. coli* identification method |  |  |  |
| --- | --- | --- | --- | --- | --- |
|  |  | Morphology<br>*E. coli* | Biochemistry<br>*E. coli* | PCR<br>(*uidA*+) | MALDI-TOF<br>*E. coli* |
| CORIANDER | 32 | 83 | 38 | 34 | 32 |
| LETTUCE | 32 | 83 | 27 | 26 | 25 |
| EV1 | 32 | 84 | 10 | 9 | 6 |
| EV2 | 32 | 84 | 3 | 2 | 1 |
| TOTAL | 128 | 334 | 78 | 71 | 64 |

Green salad 1(EV1): cucumber, lettuce, carrot; green salad 2 (EV2): cucumber, carrot, lettuce and beetroot; *uidA*<sup>+</sup>: positive for $\beta$ -glucuronidase gene

These findings highlight the significant prevalence of *E. coli* in coriander and lettuce, with a notably lower occurrence in mixed salads. The variation in contamination rates among different vegetable products underscores the importance of targeted microbial assessments and interventions to ensure food safety in diverse market settings.

### 3.2 Molecular analysis

Molecular identification of the 78 presumptive *E. coli* isolates confirmed 71 by amplifying the *uid*A+ gene (91.03%) (Fig. 1), indicating that 7 isolates were incorrectly identified as *E. coli*. Further identification using MALDI/TOF-TOF corroborated the presence of other microorganisms within the coliform group, which had biochemical characteristics like those of *E. coli*. The identified microorganisms were *Klebsiella pneumoniae* (5 isolates) and *Enterobacter cloacae* (2 isolates) and consequently were ruled out. With this additional analysis, we confirmed that out of the seventy-eight presumptive isolates, sixty-four were indeed *E. coli* (Table 1).

**Figure 1.**
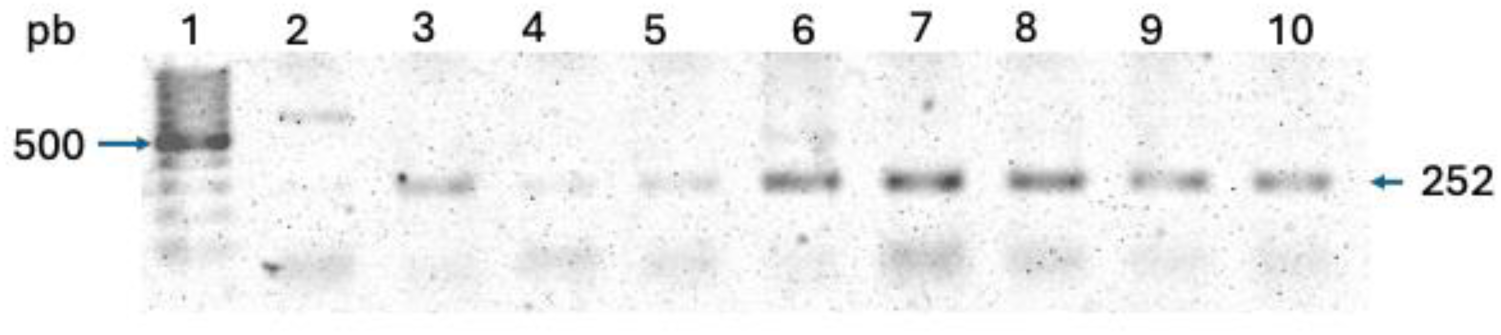
Gel electrophoresis is used to identify *Escherichia coli* using the presence of the *uid*A gene for isolates from fresh salads, lettuce, and coriander. Lane 1, 100 pb MW marker; Lane 2, negative control; Lane 3, 10 confirmed isolates.

#### 3.2.1 *E. coli* pathotype search

For the molecular identification of *E. coli* pathotypes, target genes were amplified in the 64 confirmed isolates. Of these, 51.56% were identified as enterotoxigenic *E. coli* (ETEC). We did not detect the presence of EHEC, EAEC, and EPEC pathotypes. The ETEC pathotype was determined by amplifying the *st* and *lt* genes (Fig. 1). Specifically, in 33.33% of the isolates, we successfully amplified the *lt* gene fragment. In 54.5%, we amplified only the *st* gene fragment. Both genes were found in 12.2% of the cases.

#### 3.2.2 Antimicrobial susceptibility

Most isolated ETEC resisted at least two antibiotics. Predominant resistance was found for gentamicin (97%) and cefotaxime (100%). Less resistance was observed for ampicillin (50%), sulfamethoxazole/trimethoprim, and chloramphenicol, which have a lower incidence of resistance (Table 3).

Conversely, the ETEC strains exhibited greater sensitivity to amikacin (90%) and chloramphenicol (83%). Notably, three strains demonstrated resistance to both amikacin and gentamicin. Two strains were sensitive to amphenicols (chloramphenicol) and sulphonamides, while another was sensitive to amphenicols and β-lactams.

Multidrug resistance was evaluated according to the ReLAVRA protocols (13). Table 4 shows the resistance profile of the 30 ETEC isolates evaluated with the seven antibiotics. Notably, two strains isolated from coriander exhibited extended resistance to six of seven tested antibiotics, remaining sensitive to amikacin. Among the multiresistant strains of five antibiotics, the sensitive, the sensitivity incidence was lower for amikacin (37.5%), chloramphenicol (25%), sulfamethoxazole/trimethoprim (25%), and ampicillin (17.5%) (Table 3).

## 4 Discussion

Various outbreaks of food-borne diseases have been associated with consuming contaminated fresh vegetables. Information regarding this issue is continuously updated in Europe, North America, Australia, and New Zealand due to a regulated epidemiological vigilance system. However, in developing countries in Latin America, the need for more data regarding food safety, the difficulty accessing clean water, scarce agricultural infrastructure, and limitations in implementing good farming practices present persistent challenges that prevent adequate control of gastrointestinal illnesses (14).

Vegetables can get contaminated during farming, collection, commercialization, and consumption. Vegetables consumed raw without adequate sanitization can potentially cause gastrointestinal illnesses. This highlights the importance of ensuring food is not contaminated with microorganisms, their toxins, or any other physical or chemical pollutant. Crops are often watered with wastewater, and farming soil is usually fertilized with untreated organic compost (15–18). In this work, we found the presence of ETEC in 78.78% of the coriander samples, with 21.21% of them amplifying the *lt* gene, 51.51% the *st* gene, and 6.06% both genes.

For lettuce samples, ETEC presence was detected in 9.09% of them, with 33.33% of the strains showing amplification of the lt gene, 33.33% of the *st* gene, and 33.33% of both genes (*lt/st*). Currently, there is no data on this issue in Mexico City. ETEC pathotype presence has been reported in the country on alfalfa sprouts (2%) (18); beetroot juice (2%) (19), carrot juice (1.4%) (20); round and “guaje” tomato varieties (3%-4%) (21); on a dairy product (1.78%) (22), and coriander (2%) (23).

Diarrheic sickness caused by ETEC has been reported in South Asia (27%), Latin America (22%), and Africa (North and East) 12% (24). In Bangladesh, diarrheagenic *E. coli* was responsible for 34% of diarrheic cases in adults and 90% in infants, ETEC being the prevalent strain in 23% of the samples. Likewise, ETEC and EPEC have been reported in surface waters (10, 25). Different types of diarrheagenic *E. coli* have been identified in Japan as the most frequent enteropathogenic agent, with 10% in Iran and 11% in Finland. In Thailand and Vietnam, ETEC was the pathotype causing diarrhea in 5.8% and 4% of the reported cases, respectively (10, 26, 27).

Regarding the presence of the *lt* (487 bp) and *st* (322 bp) genes, Table 6 shows that in 4 of the strains (12.12%), both gene fragments (*st*/*lt*) were amplified in 11 strains (33.3%) only the *lt* gene was amplified, and in 18 strains (54.6%) only the *st* gene was amplified. So far, no data related to the presence of both target genes on ETEC in Mexico City has been obtained. Only the *st* gene was amplified in strains originating from unpasteurized carrot juice (20) and beetroot juice (19).

The importance of the presence of both genes on ETEC lies in the fact that the *lt* toxin activates the secretion of Cl^-^ mediated by CFTR (cystic fibrosis transmembrane conductance regulator), increasing cAMP. In contrast, the *st* toxin inhibits NHE3 activity, deactivating the trafficking of the transporter (28, 29). A combination of both toxins could be crucial for developing traveler’s diarrhea. Notably, 10-14% of patients with traveler’s diarrhea after traveling to Latin America, Africa, or Asia show irritable colon symptoms (30).

Regarding antibiotic sensitivity, we found that 100% of the isolated strains were resistant to cefotaxime and 97% to gentamicin. Four strains showed resistance to β-lactams, aminoglycosides, amphenicols, and sulphonamides; nine strains displayed multi-resistance to β-lactams, amphenicols, and aminoglycosides; one strain exhibited resistance to β-lactams, amphenicols, and sulphonamides, and 17 strains had heterogeneous resistance.

This research showed the specific sensitivity of ETECs to some antibiotics: 90% for amikacin (27/30), 60% for ceftriaxone, and 3.3% for chloramphenicol. Two strains were considered sensitive to amphenicols and sulphonamides, and one exhibited sensitivity to amphenicols and β-lactams. This last data is comparable to a review from the African Continent reporting *E. coli* with 53% sensitivity to gentamicin, 49.3% to ciprofloxacin, and 47.2% to cotrimoxazole (sulphamethoxazole/trimethoprim) (31). A 2016 report found ETEC strains with multi-resistance to amikacin, gentamicin, and sulfamethoxazole/trimethoprim (32). In 2018, multi-resistance to amikacin, erythromycin, gentamicin, colistin, kanamycin, ceftriaxone, and compounds such as amoxicillin/clavulanic acid or sulphamethoxazole/trimethoprim, among others, was reported in fresh cheese (33). Comparatively, in our study, two ETEC strains showed multi-resistance to six different antibiotics, including ampicillin and gentamicin, but were sensitive to amikacin.

No published information addresses the predominance of genes (*st* or *lt*), suggesting a gene-resistance correlation. However, our results indicate that from the strains resistant to four groups of antibiotics and isolated from coriander samples, 75% showed expression of the *st* gene during molecular identification. In contrast, only 25% expressed the *lt* gene. Similarly, from the strains isolated from coriander and showing resistance to three groups of antibiotics, 71.5% expressed the *st* gene, and only 14.3% expressed the *lt* gene.

In this study, 4 of the 30 ETEC strains showed antimicrobial resistance to all the antibiotics tested (β-lactams, aminoglycosides, amphenicols, and sulphonamides). Two studies focused on multi-resistance reported *E. coli* resistance to different antibiotics. The first mentions resistance to erythromycin, amoxicillin, tetracycline, cotrimoxazole, and chloramphenicol, among others. In 2019, this microorganism resisted gentamicin, cefuroxime, cefotaxime, carbapenems and quinolones. These two studies were conducted on uropathogenic *E. coli* (UPEC) (34, 35).

Similarly, in our study, multiresistance to gentamicin, cefuroxime, cefotaxime, amphenicols (chloramphenicol), and sulphonamides (sulfamethoxazole/trimethoprim) was related to the gastrointestinal tract. Additionally, 17 strains showed heterogeneous resistance to two antibiotics of any group. Our data suggest no direct relationship between the ETEC strains and the vegetables. Thus, a direct relationship between vegetable presence, pathogen presence, and multi-resistance cannot be confirmed, posing a threat to the consumeŕs health.

Lifestyle modifications have significantly changed the consumption of ready-to-eat foods such as salads, increasing market demand. The presence of enterotoxigenic *E. coli* in fresh salads, lettuce, and coriander sold and consumed in markets or on the streets in some areas of Mexico City suggests that this bacterium could be linked to the increasing number of gastrointestinal diseases.

This study shows that enterotoxigenic *E. coli* (ETEC) strains in the analyzed vegetable products contain genes coding for toxins and exhibit antibiotic resistance, proving their pathogenic capability. Their presence on contaminated RTE salads or in vegetables such as lettuce and coriander are an actual risk for consumers.

## 5 Recommendations and future research

Vegetables can get contaminated during farming, collection, commercialization, and consumption. This means that vegetables consumed raw without proper sanitization can cause gastrointestinal illnesses, highlighting the importance of ensuring that our food is not contaminated with microorganisms or toxins or even with physical or chemical pollutants. This problem is unsurprising because crops are often watered with wastewater, and the soil is fertilized with untreated organic compost. Additionally, coriander, a prevalent ingredient in dishes or salsas in Mexican cuisine, poses a security risk to consumers due to the lack of hygiene and sanitization applied to its handling. It is common for people to store it in buckets filled with non-drinking water all day long, making proper handling and food preparation critical for safety.

Consuming food prepared and sold on the streets or in open markets is risky, as it can contain harmful microorganisms, posing a substantial health risk to consumers of all ages and social classes. The presence of enterotoxigenic *E. coli* in fresh salads, lettuce, and coriander sold or consumed in markets or the streets in some areas of Mexico City, combined with lifestyle changes that have generated the demand for vegetables, suggests that this bacterium could be related to the increasing number of gastrointestinal diseases.

## 6 Material and Methods

### 6.1 Sampling

Between January 2018 and February 2020, 128 samples of vegetable products were analyzed. These samples included 32 samples of lettuce, cucumber, and carrot salads obtained from the “La Vicentina” market (EV1); 32 samples of cucumber, lettuce, carrot, and beet salads obtained from the “La Purísima” market (EV2); 32 samples of romaine lettuce and 32 samples of coriander from the “Central de Abastos” market. All collection sites were in the Iztapalapa municipality in Mexico City. Each sample weighed approximately 300 g and was transported in a hermetically sealed container at an approximate temperature of 10°C. The time elapsed between collection and analysis was kept to under an hour to ensure the integrity and accuracy of the results.

### 6.2 Microbiological analysis

For *E. coli* isolation and identification, the methodology recommended by the FDA’s Bacteriological Analytical Manual (36) and the NOM 210-SSa1-2014 were followed. All the sample in its original container was chopped into small pieces. Afterward, 10 grams were placed on 90 ml of 0.1% peptone water (Difco®) for a 10-1 dilution. A series of ten-fold dilutions were prepared up to 10^-3^. For the MPN of fecal coliforms, a 3.3.3 series was used. 1 ml of each dilution was placed on tubes containing 9 ml lactose broth with an inverted Durham tube. The tubes were incubated at 35 ± 2° C during 48 ± 2 h. The tubes showing growth and gas production were inoculated by transferring 2-3 loops of inoculum onto *E. coli* broth (EC) (Difco®) containing inverted Durham’s tube. They were incubated at 45.5 ± 0.2° C in an agitated water bath for 48 h. Biochemical tests for the identification of *E. coli* were performed.

### 6.3 Molecular analysis

#### 6.3.1 Bacterial DNA isolation and PCR

Presumptive *E. coli* was inoculated on capped test tubes containing 5 mL of Brain Heart Infusion broth (BHI) and incubated at 37°C for 24 h. Wizard ® Genomic DNA purification (PROMEGA) commercial kit was used following the manufacturer-recommended protocol.

#### 6.3.2 Amplification of the target genes

PCR reaction was standardized for the amplification of the fragments of the target *E. coli* genes and for every pathotype. Reaction mix consisted of 12.5 µL Master mix 2X (Thermo Scientific®), 6.5 µL of nuclease free water, 2 µL of forward primer, 2μL of reverse primer (Table 1 supplementary material) and 2 μL of DNA in a total volume of 25µL.

Conditions used for the *uid*A gene amplification were denaturalization at 94°C/5 min followed by 30 cycles at 55°C/30 s, 72°C/1 min, and a final extension at 72°C/7 min. Amplification conditions for the *pet* and *aggR* (EAEC) genes and ETEĆs *It* and *st* genes included an initial denaturalization step at 94°C/5 min, followed by 30 cycles at 94°C/ 1min, 52°C/1 min, 72°C/1 min, and a final extension at 72°/7min. For the *stx1* and *stx2* (EHEC) genes and *eaeA* and *bfpA* (EPEC) genes, an initial denaturalization was performed at 94°C/5 min followed by 30 cycles at 94°C/ 1min, 58°C/1min, 72°C/1min and a final extension at 72°C/7min.

A Labnet MultiGene^tm^ Gradient PCR Thermal Cycler (TC-9600G-230V, Staffordshire, UK) was used, and PCR amplification products were visualized by electrophoresis on a 1.5% agarose gel where 5 μL of each sample were mixed with 2 μL of Green loading buffer (Jenna Bioscience®). Electrophoresis was performed using Tris-Glacial acetic acid-EDTA (TAE) buffer applying 60 V current for 30 minutes. 100 bp molecular weight was used. Finally, the amplification products were visualized on a blue light transilluminator Model MBE-150 Major Science®. *E. coli* EAEC ATCC29552, EHEC ATCC 43895, EPEC ATCC 43887, and ETEC ATCC 35401 strains were used as positive controls and were donated by the Molecular Microbiology Laboratory of the Escuela Nacional de Ciencias Biológicas. The strains were maintained at -20°C in BHI broth +10% glycerol until their use.

### 6.4 Strain identification confirmation with the MALDI Biotyper® System

*E. coli* confirmed isolates using biochemical identification were analyzed by the MALDI Biotyper protocol from Bruker Daltonics on a mass spectrophotometer Autoflex Speed MALDI-TOF/TOF, using an α-Cyano-4-hydroxycinnamic acid matrix in 50% acetonitrile, 2.5% trifluoroacetic acid and Milli-Q® water to a final concentration of 10 mg/mL.

Each sample was processed as follows: 300 µL of Milli-Q water + culture was mixed with 900 µL of ethanol and centrifuged at 14,000 rpm on an Eppendorf centrifuge, and the supernatant was discarded. The pellets were left to dry and resuspended in 5, 10, or more μL of 70% formic acid, depending on the sample amount. This mixture was homogenized with a micropipette, and the same volume of acetonitrile was added and centrifuged one more time. Afterward, 1µL of the supernatant was placed on each well of a 98-sample stainless steel MALDI plate containing 1 μl of the matrix. Manufacturer directions were followed for the interpretation of the results.

### 6.5 Antimicrobial susceptibility

The methodology described by the CLSI (Clinical & Laboratory Standards Institute) was followed for antimicrobial susceptibility. *E. coli* strains were inoculated on tubes containing 5 mL of trypticasein soy broth and incubated at 37°C for 24 h. Afterward, 100 μl of these pre-cultures were inoculated on tubes containing 5 mL of trypticase soy broth. They were incubated with agitation for 3 hours until they reached turbidity like the MacFarland 0.5 nephelometer tube.

These cultures were streaked massively onto a Mueller-Hinton agar plate where 7 filter paper discs were placed on equidistant zones, each one impregnated with the following antibiotics: Amikacin (30 µg), Ampicillin (30 µg), Cefotaxime (30 µg), Ceftriaxone (30 µg), Chloramphenicol (30 µg), Gentamicin (10 µg) and Sulfamethoxazole/Trimethoprim (25 µg). Plaques were incubated at 37°C for 24, and afterward, the inhibition zones were measured as determined by the CLSI on M100 ED31:2021. Each assay was performed in duplicate to ensure the precision of the results.

The antibiotic susceptibility assay results were compared with the zone diameter table for Enterobacterales of the M100 ED31:2021 data from the CLSI (Table 2 supplementary material) (37).

**Table 2.** Antimicrobial activity incidence shown by different ETEC strains isolated from several vegetables collected in Mexico City.

| Antibiotic | Resistance |  |  |  |  | Sensitivity |  |  |  |  |
| --- | --- | --- | --- | --- | --- | --- | --- | --- | --- | --- |
|  | Lettuce | Coriander | EV1 | EV2 | Total | Lettuce | Coriander | EV1 | EV2 | Total |
| AK | 0 | 2 | 1 | 0 | 3 | 2 | 22 | 1 | 2 | 27 |
| AM | 1 | 10 | 2 | 1 | 14 | 1 | 14 | 0 | 0 | 15 |
| CFX | 2 | 24 | 2 | 2 | 30 | 0 | 0 | 0 | 0 | 0 |
| CTX | 1 | 9 | 1 | 1 | 12 | 1 | 15 | 1 | 1 | 18 |
| CL | 0 | 4 | 0 | 1 | 5 | 2 | 20 | 2 | 1 | 25 |
| GE | 2 | 23 | 2 | 2 | 29 | 0 | 1 | 0 | 0 | 1 |
| STX | 0 | 12 | 1 | 0 | 13 | 2 | 12 | 1 | 2 | 17 |

**Table 3.**
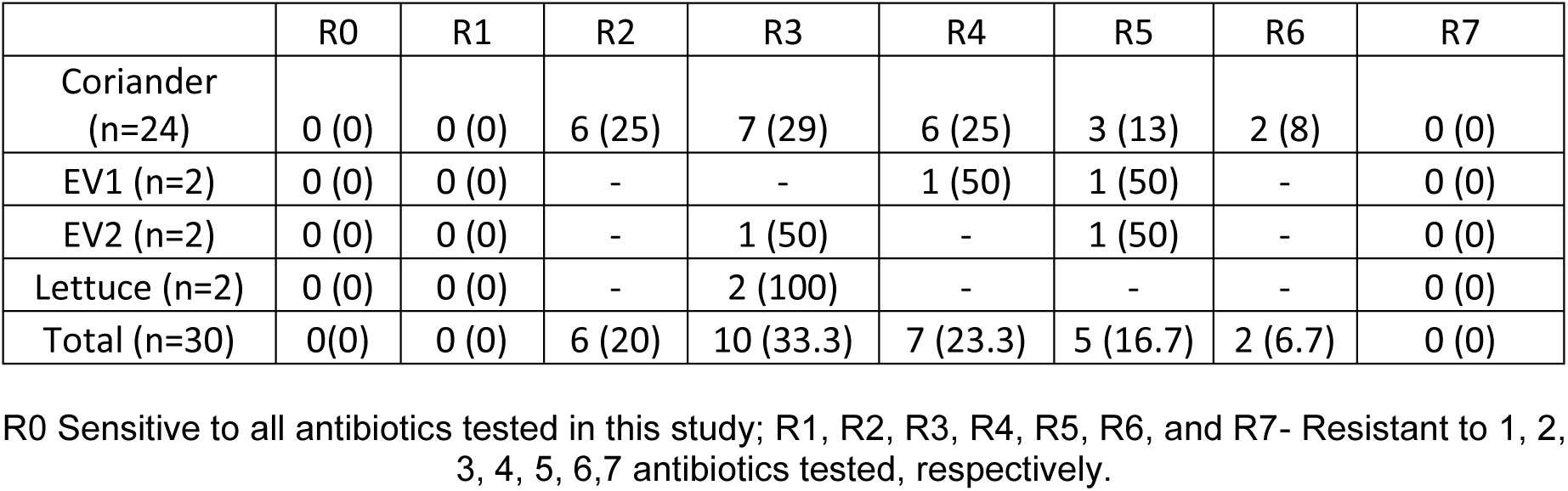
Multiresistance in identified ETEC strains isolated from several vegetables in Mexico City.

|  | R0 | R1 | R2 | R3 | R4 | R5 | R6 | R7 |
| --- | --- | --- | --- | --- | --- | --- | --- | --- |
| Coriander (n=24) | 0 (0) | 0 (0) | 6 (25) | 7 (29) | 6 (25) | 3 (13) | 2 (8) | 0 (0) |
| EV1 (n=2) | 0 (0) | 0 (0) | - | - | 1 (50) | 1 (50) | - | 0 (0) |
| EV2 (n=2) | 0 (0) | 0 (0) | - | 1 (50) | - | 1 (50) | - | 0 (0) |
| Lettuce (n=2) | 0 (0) | 0 (0) | - | 2 (100) | - | - | - | 0 (0) |
| Total (n=30) | 0(0) | 0 (0) | 6 (20) | 10 (33.3) | 7 (23.3) | 5 (16.7) | 2 (6.7) | 0 (0) |

R0 Sensitive to all antibiotics tested in this study; R1, R2, R3, R4, R5, R6, and R7- Resistant to 1, 2, 3, 4, 5, 6,7 antibiotics tested, respectively.

## 7 Data availability

The data used are included in this publication’s supplemental material. The corresponding author can send all remaining data and strains upon request.

## 9 Acknowledgments

Funding Universidad Autónoma Metropolitana, No. 5/65.21, Instituto Politécnico Nacional, ENCB No. 20241508 y Conahcyt fellowship to CRVQ no.236826.

## 10 Author contributions

Vázquez-Quiñones, C. R. contribution to the concept or design of the article; collected the data, performed the analysis, interpreted the data, wrote the paper, Rincón-Guevara, M.A. The acquisition, analysis, or interpretation of data Approved the version to be published.

Natividad-Bonifacio, I. Contribution to the concept or design of the article, interpretation of data for the article, revised critically of content, approved the version to be published, funding.

Vázquez-Salinas, C. Contributed to the concept or design of the article, interpreted data for the article, revised content critically, approved the version to be published, and funded.

González-Márquez, H. Contribution to the concept or design of the article, interpretation of data for the article, critically revised content, approval of the version to be published, and funding.

## SUPPLEMENTARY MATERIALS

**Table S1.**
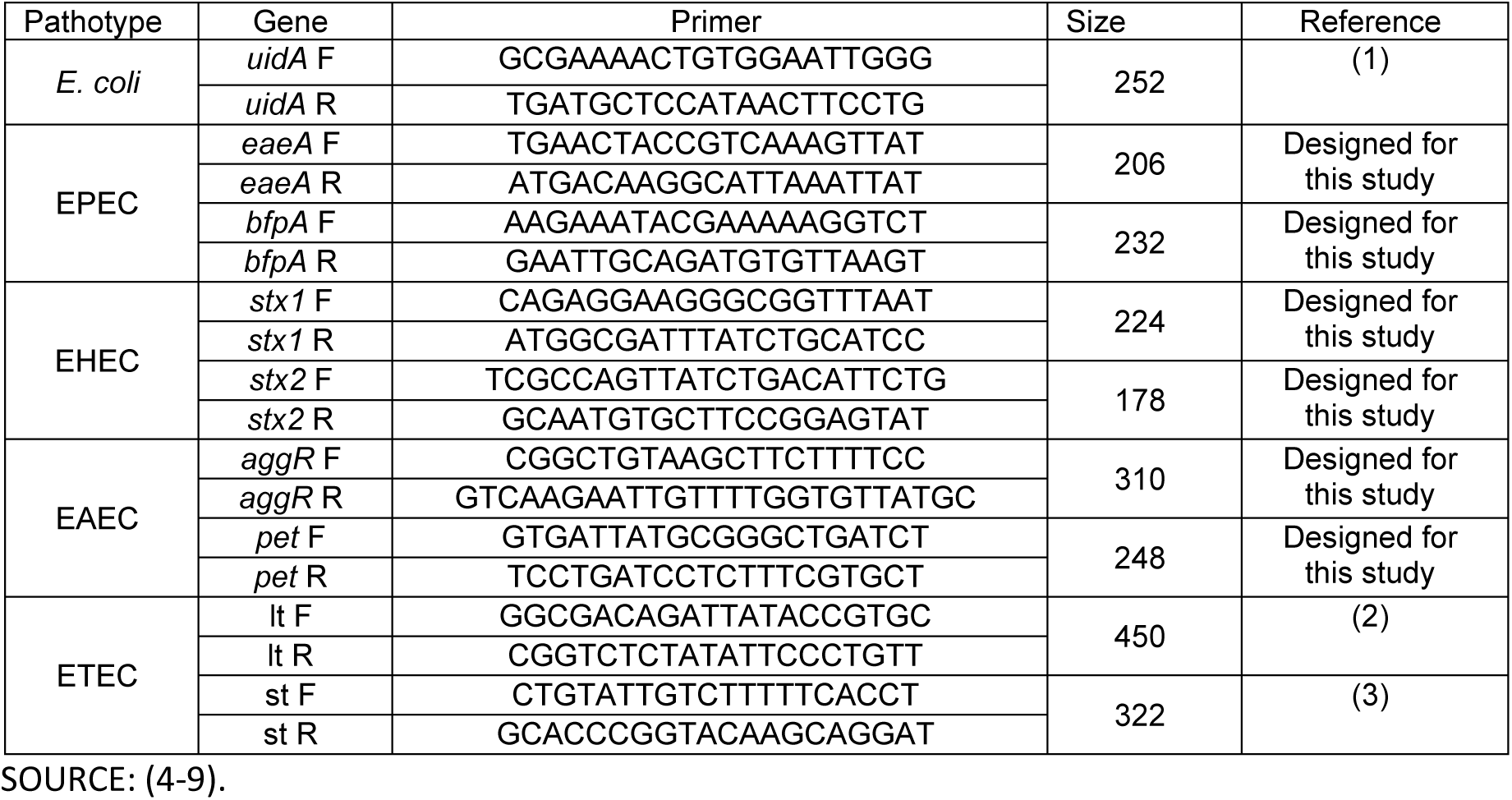
Primer sequence for the target genes of different *Escherichia coli* pathotypes, amplicon size, and Tm referred to in other references.

**Table S2.** Antibiotic data and inhibition zone diameter for Enterobacterales were used to determine the ETEC strains’ sensitivity.

| Antibiotic | Abbreviation | Units (µg) | Sensitive | Intermediate | Resistant |
| --- | --- | --- | --- | --- | --- |
| Amikacin | AK | 30 | ≥ 17 | 15-16 | ≤ 14 |
| Ampicillin | AM | 30 | ≥ 17 | 14-16 | ≤ 13 |
| Cefotaxime | CFX | 30g | ≥ 26 | 23-25 | ≤ 22 |
| Ceftriaxone | CTX | 30 | ≥ 23 | 20-22 | ≤ 19 |
| Chloramphenicol | CL | 30 | ≥ 18 | 13-17 | ≤ 12 |
| Gentamicin | GE | 10 | ≥ 15 | 13-14 | ≤ 12 |
| Sulfamethoxazole/<br>Trimethoprim | STX | 25 | ≥ 16 | 11-159 | ≤ 10 |

Measure units in the inhibition zones are mm.

**Table S3.** MALDI-TOF/TOF confirmation of ETEC isolates in vegetables and salads.

| **Isolate** | **Date/sample** | **Origin** | **Biochemical confirmation** | **MALDI TOF identification** |
| --- | --- | --- | --- | --- |
| 1 | MAY 18 | Fresh salad 1 | *Escherichia coli* | *Escherichia coli* |
| 2 | MAY 18 | Fresh salad 1 | *Escherichia coli* | *Escherichia coli* |
| 3 | MAY 18 | Lettuce | *Escherichia coli* | *Enterobacter cloacae* |
| 4 | MAY 18 | Lettuce | *Escherichia coli* | *Escherichia coli* |
| 5 | MAY 18 | Fresh salad 2 | *Escherichia coli* | *Escherichia coli* |
| 6 | AUG 18 | Fresh salad 1 | *Escherichia coli* | *Escherichia coli* |
| 7 | AUG 18 | Coriander | *Escherichia coli* | *Klebsiella Pneumoniae* |
| 8 | SEP 18 | Coriander | *Escherichia coli* | *Escherichia coli* |
| 9 | SEP 18 | Coriander | *Escherichia coli* | *Escherichia coli* |
| 10 | SEP 18 | Coriander | *Escherichia coli* | *Escherichia coli* |
| 11 | SEP 18 | Coriander | *Escherichia coli* | *Escherichia coli* |
| 12 | SEP 18 | Coriander | *Escherichia coli* | *Escherichia coli* |
| 13 | SEP 18 | Coriander | *Escherichia coli* | *Escherichia coli* |
| 14 | SEP 18 | Coriander | *Escherichia coli* | *Escherichia coli* |
| 15 | SEP 18 | Coriander | *Escherichia coli* | *Escherichia coli* |
| 16 | SEP 18 | Coriander | *Escherichia coli* | *Escherichia coli* |
| 17 | SEP 18 | Coriander | *Escherichia coli* | *Escherichia coli* |
| 18 | SEP 18 | Coriander | *Escherichia coli* | *Escherichia coli* |
| 19 | SEP 18 | Coriander | *Escherichia coli* | *Escherichia coli* |
| 20 | SEP 18 | Coriander | *Escherichia coli* | *Escherichia coli* |
| 21 | SEP 18 | Coriander | *Escherichia coli* | *Escherichia coli* |
| 22 | SEP 18 | Coriander | *Escherichia coli* | *Escherichia coli* |
| 23 | SEP 18 | Lettuce | *Escherichia coli* | *Escherichia coli* |
| 24 | SEP 18 | Lettuce | *Escherichia coli* | *Escherichia coli* |
| 25 | SEP 18 | Lettuce | *Escherichia coli* | *Escherichia coli* |
| 26 | SEP 18 | Lettuce | *Escherichia coli* | *Escherichia coli* |
| 27 | SEP 18 | Lettuce | *Escherichia coli* | *Escherichia coli* |
| 28 | SEP 18 | Lettuce | *Escherichia coli* | *Escherichia coli* |
| 29 | SEP 18 | Lettuce | *Escherichia coli* | *Escherichia coli* |
| 30 | SEP 18 | Lettuce | *Escherichia coli* | *Escherichia coli* |
| 31 | SEP 18 | Lettuce | *Escherichia coli* | *Escherichia coli* |
| 32 | SEP 18 | Fresh salad 1 | *Escherichia coli* | *Escherichia coli* |
| 33 | SEP 18 | Fresh salad 1 | *Escherichia coli* | *Klebsiella Pneumoniae* |
| 34 | SEP 18 | Coriander | *Escherichia coli* | *Escherichia coli* |
| 35 | SEP 18 | Coriander | *Escherichia coli* | *Enterobacter cloacae* |
| 36 | SEP 18 | Coriander | *Escherichia coli* | *Escherichia coli* |
| 37 | SEP 18 | Coriander | *Escherichia coli* | *Escherichia coli* |
| 38 | SEP 18 | Coriander | *Escherichia coli* | *Escherichia coli* |
| 39 | SEP 18 | Coriander | *Escherichia coli* | *Escherichia coli* |
| 40 | SEP 18 | Lettuce | *Escherichia coli* | *Escherichia coli* |
| 41 | SEP 18 | Lettuce | *Escherichia coli* | *Escherichia coli* |
| 42 | SEP 18 | Lettuce | *Escherichia coli* | *Escherichia coli* |
| 43 | SEP 18 | Lettuce | *Escherichia coli* | *Escherichia coli* |
| 44 | SEP 18 | Lettuce | *Escherichia coli* | *Escherichia coli* |
| 45 | SEP 18 | Lettuce | *Escherichia coli* | *Escherichia coli* |
| 46 | SEP 18 | Lettuce | *Escherichia coli* | *Escherichia coli* |
| 47 | SEP 18 | Lettuce | *Escherichia coli* | *Escherichia coli* |
| 48 | SEP 18 | Lettuce | *Escherichia coli* | *Escherichia coli* |
| 49 | SEP 18 | Lettuce | *Escherichia coli* | *Escherichia coli* |
| 50 | SEP 18 | Lettuce | *Escherichia coli* | *Escherichia coli* |
| 51 | SEP 18 | Lettuce | *Escherichia coli* | *Escherichia coli* |
| 52 | OCT 18 | Coriander | *Escherichia coli* | *Escherichia coli* |
| 53 | OCT 18 | Coriander | *Escherichia coli* | *Klebsiella Pneumoniae* |
| 54 | OCT 18 | Lettuce | *Escherichia coli* | *Klebsiella Pneumoniae* |
| 55 | OCT 18 | Lettuce | *Escherichia coli* | *Escherichia coli* |
| 56 | OCT 18 | Lettuce | *Escherichia coli* | *Klebsiella Pneumoniae* |
| 57 | OCT 18 | Fresh salad 1 | *Escherichia coli* | *Klebsiella Pneumoniae* |
| 58 | OCT 18 | Fresh salad 1 | *Escherichia coli* | *Escherichia coli* |
| 59 | OCT 18 | Fresh salad 1 | *Escherichia coli* | *Escherichia coli* |
| 60 | OCT 18 | Fresh salad 1 | *Escherichia coli* | *Escherichia coli* |
| 61 | OCT 18 | Fresh salad 1 | *Escherichia coli* | *Escherichia coli* |
| 62 | JUL 19 | Coriander | *Escherichia coli* | *Escherichia coli* |
| 63 | JUL 19 | Coriander | *Escherichia coli* | *Escherichia coli* |
| 64 | JUL 19 | Coriander | *Escherichia coli* | *Escherichia coli* |
| 65 | JUL 19 | Coriander | *Escherichia coli* | *Escherichia coli* |
| 66 | JUL 19 | Lettuce | *Escherichia coli* | *Escherichia coli* |
| 67 | JUL 19 | Fresh salad 2 | *Escherichia coli* | *Klebsiella Pneumoniae* |
| 68 | JUL 19 | Fresh salad 2 | *Escherichia coli* | *Escherichia coli* |
| 69 | AGO 19 | Coriander | *Escherichia coli* | *Escherichia coli* |
| 70 | AGO 19 | Coriander | *Escherichia coli* | *Escherichia coli* |
| 71 | AGO 19 | Coriander | *Escherichia coli* | *Escherichia coli* |
| 72 | AGO 19 | Coriander | *Escherichia coli* | *Escherichia coli* |
| 73 | AGO 19 | Coriander | *Escherichia coli* | *Escherichia coli* |
| 74 | OCT 19 | Coriander | *Escherichia coli* | *Escherichia coli* |
| 75 | OCT 19 | Coriander | *Escherichia coli* | *Escherichia coli* |
| 76 | OCT 19 | Coriander | *Escherichia coli* | *Escherichia coli* |
| 77 | OCT 19 | Coriander | *Escherichia coli* | *Escherichia coli* |
| 78 | OCT 19 | Coriander | *Escherichia coli* | *Escherichia coli* |

**Table 4.**
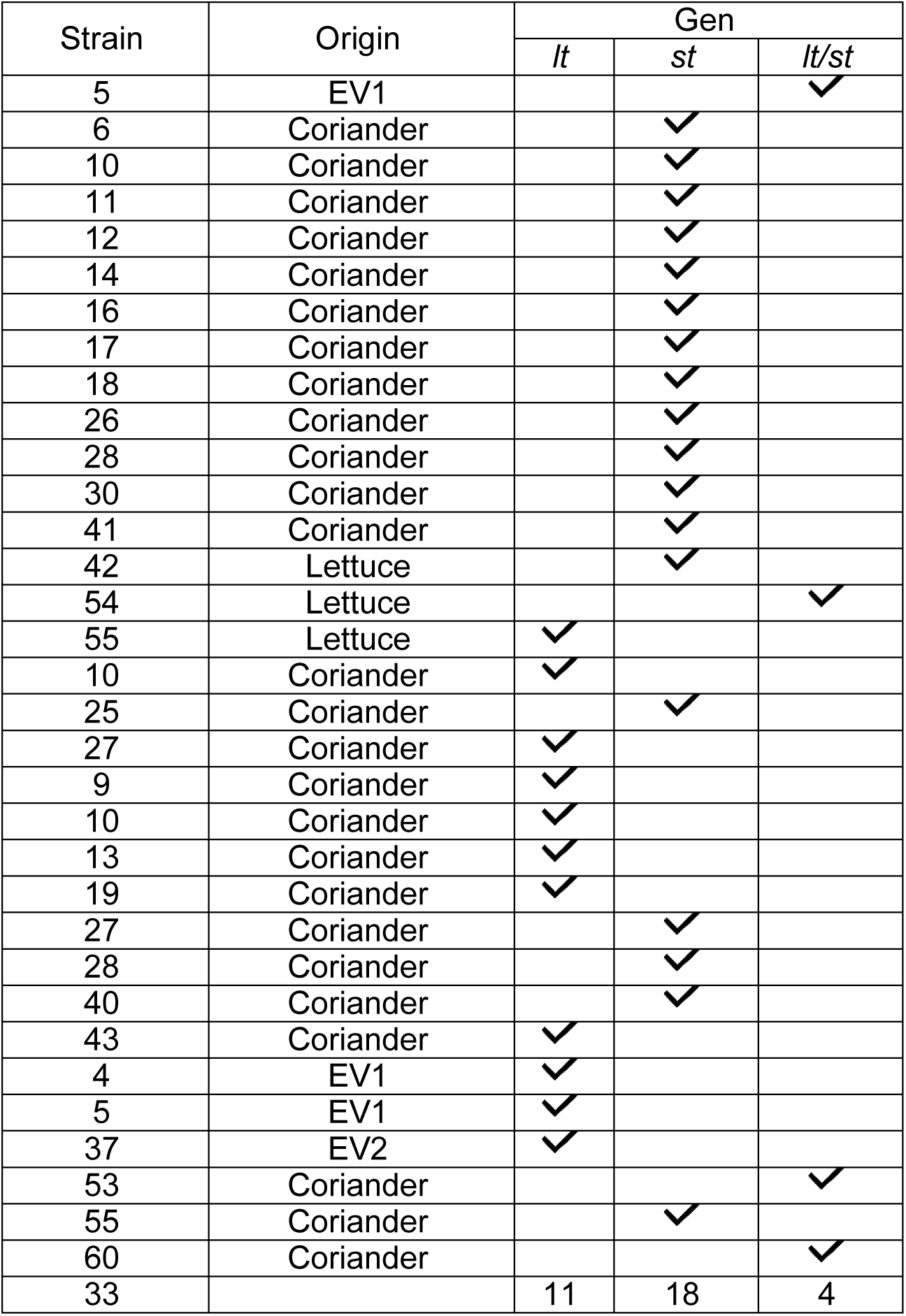
Presence of *lt* and *st* genes in confirmed *E. coli* strains isolated from fresh salads collected in CDMX (Iztapalapa) between May-October 2018.

| Strain | Origin | Gen |  |  |
| --- | --- | --- | --- | --- |
|  |  | *lt* | *st* | *lt/st* |
| 5 | EV1 |  |  | ✓ |
| 6 | Coriander |  | ✓ |  |
| 10 | Coriander |  | ✓ |  |
| 11 | Coriander |  | ✓ |  |
| 12 | Coriander |  | ✓ |  |
| 14 | Coriander |  | ✓ |  |
| 16 | Coriander |  | ✓ |  |
| 17 | Coriander |  | ✓ |  |
| 18 | Coriander |  | ✓ |  |
| 26 | Coriander |  | ✓ |  |
| 28 | Coriander |  | ✓ |  |
| 30 | Coriander |  | ✓ |  |
| 41 | Coriander |  | ✓ |  |
| 42 | Lettuce |  | ✓ |  |
| 54 | Lettuce |  |  | ✓ |
| 55 | Lettuce | ✓ |  |  |
| 10 | Coriander | ✓ |  |  |
| 25 | Coriander |  | ✓ |  |
| 27 | Coriander | ✓ |  |  |
| 9 | Coriander | ✓ |  |  |
| 10 | Coriander | ✓ |  |  |
| 13 | Coriander | ✓ |  |  |
| 19 | Coriander | ✓ |  |  |
| 27 | Coriander |  | ✓ |  |
| 28 | Coriander |  | ✓ |  |
| 40 | Coriander |  | ✓ |  |
| 43 | Coriander | ✓ |  |  |
| 4 | EV1 | ✓ |  |  |
| 5 | EV1 | ✓ |  |  |
| 37 | EV2 | ✓ |  |  |
| 53 | Coriander |  |  | ✓ |
| 55 | Coriander |  | ✓ |  |
| 60 | Coriander |  |  | ✓ |
| 33 |  | 11 | 18 | 4 |

